# Unravelling the anticancer and enzyme inhibition potential of different classes of compounds including a new steroid, isolated from *Cassia mimosoïdes*

**DOI:** 10.1101/2023.03.05.531233

**Authors:** Robert Viani Kepdieu Tchebou, Umar Farooq, Sara Khan, Azhar Rasool, Rizwana Sarwar, Aneela Khushal, Léon Azefack Tapondjou, Syed Majid Bukhari, Rémy Bertrand Teponno

## Abstract

The genus *Cassia* is a significant source of secondary metabolites that are physiologically active and come from several chemical classes. The current research deals with the isolation, spectroscopic elucidation (1D and 2D NMR spectroscopy) and enzymatic activity of fifteen known compounds as well as a new unidentified avenasterol derivative namely 21-methylene-24-ethylidene lophenol. The urease and *β*-glucosidase inhibitory effects of these compounds were studied for the first time, and molecular docking studies were also performed to verify the structure-activity relationships. All the compounds evaluated towards urease showed higher inhibitory activity (1.224±0.43 < IC_50_ > 6.678±0.11 *μ*M) compared to standard thiourea (IC_50_ = 18.61±0.11 *μ*M). Molecular docking results revealed that compound **7** strongly inhibits urease due to the formation of a stable ligand-urease complex via hydrogen bonding, van der Waal and hydrophobic interactions. Formation of a favourable complex of **7** with the target enzyme gave a more negative docking score (−6.95 kcal/mol) than that of thiourea (−3.13 kcal/mol). Regarding the *β*-glucosidase enzyme, all the compounds evaluated did not show activity except compound **1** which inhibited the latter with a percentage of inhibition of 82.6. These findings imply that this plant may be a contender for developing novel treatments for infectious disorders brought on by urease-producing bacteria.

## Introduction

Various medicinal plants have been used for years in daily life to treat diseases around the world. The interest in medicinal plants reflects the recognition of the validity of many traditional claims regarding the value of natural products in health care. *Cassia mimosoides* L. (Caesalpiniaceae), is distributed in various countries including Cameroon, Luzon, Mindanao, India, China, Malaysia and Australia. This plant is widely used by tribal people to treat various ailments including typhoid fever and other microbial infections (Kakpo et al., 2019; Malzy, 1954) It is used in Uganda to treat pediatric cough (Namukobe et al., 2011). In northwestern Tanzania, the aerial parts of *C. mimosoides* are pounded and mixed with animal fat, applied topically or taken orally for fractures, cleaning of the uterus by pregnant women, and as an antibacterial. In addition, some South African diviners used these plants for their oneirogenic properties (Sobiecki, 2008). The roots are used to treat diarrhea, colic, dysentery, and stomach spasms.

Parts of *Cassia mimosoides* are known to be an important source of secondary metabolites, including anthraquinones. Physcion, chrysophanol, 1,8-dihydroxy-6-methoxy-2-methyl anthraquinone, 1,8-dihydroxy-6-methoxy-3-methyl anthraquinone, and emodin have been reported from the aerial part (Dave & Ledwani, 2012). Luteolin, emodin, 1,3-benzenediol, oleanolic acid, (*R*)-artabotriol, *α*-L-rhamnose, *β*-sitosterol, and daucosterol were isolated from ethanolic extract of *Cassia mimosoides* (J.-D. Zhang et al., 2009). In addition to phenolic compounds and their derivatives, phytochemical screening of the latter revealed the presence of alkaloids, steroids, saponins, carbohydrates, tannins, glycosides, proteins, and amino acids (Prusty et al., 2011).

Ureases are common metalloenzymes made by bacteria, fungi, and plants but not by animals. They quickly catalyse the hydrolysis of urea to produce ammonia and carbamate, followed by the urea’s breakdown into a second ammonia molecule and carbon dioxide. By giving bacteria nitrogen in the form of ammonia for their growth, the enzyme plays a critical part in their pathogenicity. The pathogenicity of gastrointestinal disorders such as gastritis, duodenal, peptic ulcer, and gastric cancer is significantly influenced by *Helicobacter pylori’s* ureolytic activity. Ammonia produced by ureases is responsible for human and animal cases of hepatic encephalopathy, hepatic coma, urolithiasis, pyelonephritis, and urinary catheter encrustation. In light of this, urease inhibitors have garnered a lot of interest as potential treatments for infections brought on and facilitated by ureolytic activity. Several urease inhibitors, such as fluorofamide, hydroxyureas, and hydroxamic acids have been reported in the past. Some of these inhibitors have, however, been banned from usage *in vivo* because of their instability or toxicity. Active metabolites found in plants are well known to be helpful in treating a variety of viral disorders. In order to address concerns about toxicity right away, a lot of emphasis has been paid to examining the unique biological features of phytochemicals extracted from food plants. It has been demonstrated that a number of medicinal plants, herbs, extracts, and isolated substances have anti-urease properties. Lysosomal storage disorders (LSDs) are monogenic disorders caused by disturbances in the activity of lysosomal proteins, leading to the accumulation of undegraded metabolites (often called storage products) in the lysosome. There are more than 40 known lysosomal disorders, which can be divided into broad categories according to the nature of the accumulated substances: sphingolipidosis, oligosaccharidosis and mucopolysaccharidosis (Futerman & Meer, 2004). In most cases, sphingolipidosis or sphingolipid overload disease is due to reduced catalytic activity of one or more lysosomal hydrolases involved in one of the steps of lysosomal overload degradation and leads to intracellular accumulation of undegraded lysosomal overload (Raas-Rothschild et al., 2004). For example, Gaucher disease, one of the most common lysosomal overload diseases, is caused by mutations in the gene encoding acid beta-glucosidase, resulting in the accumulation of glucosylceramide in the lysosome. Fabry disease is the result of a defect in alpha-galactosidase activity, resulting in the accumulation of globulotrisaccharide ceramides. Inhibitors such as glucosidase are responsible for disrupting the activity of glucosidase, an enzyme that cleaves glycosidic bonds. By altering or blocking specific metabolic processes, these inhibitors play an important role in revealing the function of glucosidases in living systems. This discovery has led to a variety of applications for these chemical entities in agriculture and medicine (Asano, 2003).

Glycosidases are very important enzymes, widely used in the fields of biotechnology, food, wood chemistry or medical research. They are particularly used for the enzymatic synthesis of complex oligosaccharides under mild conditions. Numerous inhibitors of these enzymes exist and play an important role both in furthering our knowledge of the mechanisms of enzymatic hydrolysis of carbohydrates and as therapeutic tools. Among these inhibitors, iminosugars and carbasugars, analogues of sugars in which the oxygen atom has been replaced by a nitrogen atom and a carbon atom respectively, are particularly active compounds. Some of them are even currently marketed as anti-diabetics. Recently, two new glycosidase inhibitors, salacinol and kotalanol, with a unique zwitterionic structure, were isolated from *Salacia reticulata*, found in southern India and Sri Lanka1. The roots and stems of this plant are used in traditional pharmacopoeia to treat diabetes (Yoshikawa et al., 1997, 1998).

The search for novel glucosidase inhibitors is very important, as they have potential therapeutic applications in the treatment of diabetes, human immunodeficiency virus infection, metastatic cancer, and lysosomal overload diseases. These glucosidase inhibitors can also be used to explore biochemical pathways and understand the structure-activity relationship patterns required to mimic enzymatic transition states (de Melo et al., 2006). There is therefore a need to identify alternative, safe and less expensive inhibitors for the treatment of these various conditions that undermine the health of populations. Since plants are the primary source of drugs, the search for new *β*-glucosidase inhibitors can begin from the screening of plants with therapeutic properties and used by traditional healers as remedies for related opportunistic infections. The search for new drugs with a broad spectrum of action and few side effects on humans prompted us to conduct DFT and docking studies on the in-silico and in-vitro *β*-glycosidase enzymatic activity of compounds from *Cassia mimosoïdes*.

### Method: General experimental procedures

Acetone-*d_6_*, chloroform-*d*, metthanol-*d_4_*, and DMSO-*d_6_* were used as the analytical solvent for ^1^H and ^13^C NMR spectra (Bruker AMX-400 MHz and a Bruker Avance III-600 MHz). Tetramethylsilane (TMS) was used as a reference to the solvent residue while coupling constants (*J*) and chemical shifts (*δ*) are expressed in Hz and parts per million units (ppm) respectively. Silica gel 60 (0.040-0.063 mm, Merck), 60 F254 (Merck), OSD, and Sephadex LH-20 gel were used in different chromatographic techniques (HPLC, column, and TLC). Sulfuric acid 10% and UV light (254 and 365 nm) were used to revealdifferent secondary metabolites.

### Plant material

The aerial part of *C. mimosoïdes* was collected in Dschang (5°27′0′′N, 10°4′0′′E), West Region of Cameroon, in October 2018 and identified at the National Herbarium of Cameroon (NHC), Yaounde, where a voucher specimen (No. 8521/ HCN) was deposited.

### Extraction and isolation of the plant constituents

The aerial part of *C. mimosoïdes* was cut, dried and ground to give 6 Kg of powder. 4 Kg of this powder was extracted with ethanol (20 L) 3 × 24 h at room temperature followed by filtration. The filtrate obtained was concentrated under reduced pressure to give a crude extract (141 g). The extract was suspended in water and successively partitioned to ethyl acetate and butanol fractions. The ethyl acetate and *n*-butanol fractions were subjected to silica gel column chromatography using different solvent mixtures (*n*-hexane-ethyl acetate, ethyl acetate-methanol and ethyl acetate-methanol-water) in increasing polarity to give twelve subfractions (A-M). By repeating different chromatographic techniques (column and HPLC) of these twelve fractions, one new compound (**1**) and fifteen known secondary metabolites (**2-16**) were obtained (Figure 1). It should be noted, that all of these known compounds are reported herein for the first time from *Cassia mimosoïdes*.

**Fig 1.**
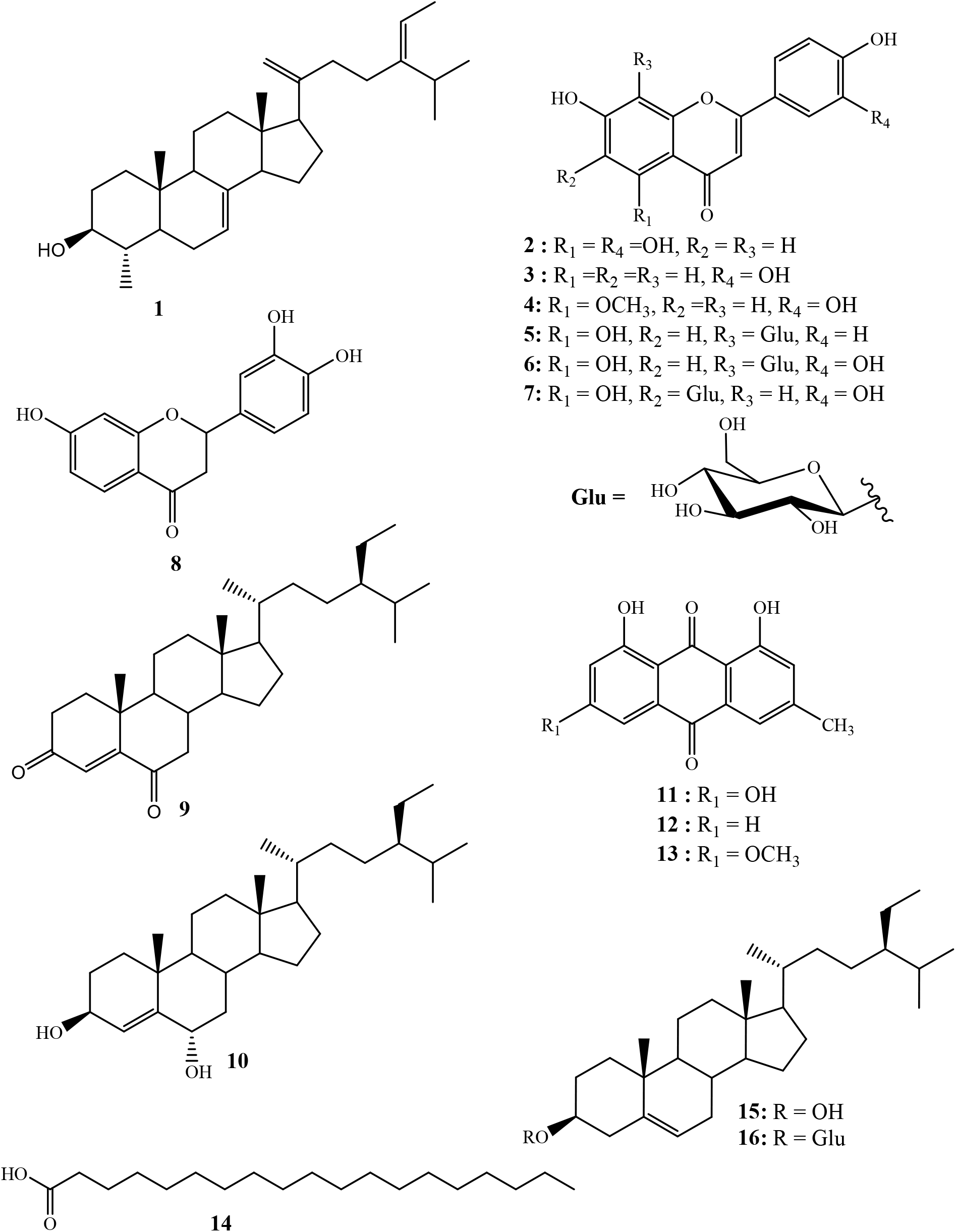
Structures of compounds (**1**-**16**) isolated from *Cassia mimosoïdes*.

#### Molecular modeling

The structure of the isolated secondary metabolites was performed using ChemDraw Professional 15.0 and their molecular docking simulations were performed using MOE (Molecular Operating Environment) software (version 2019.0102). For protein-ligand interactions, crystal structures of urease and *β*-glucosidase enzymes (PDB: 3LA4 and 2XYH) were extracted from the Protein Data Bank (PDB). Removal of water, co-crystallized ligands, and hydrogen addition were applied during the preparation process of the target proteins. Binding energy, S-score function and RMSD were used to rank the compounds. Based on their energies and interactions with the active site residues of the *β*-glucosidase protein, the highest ranked poses were selected for further study. Default settings were used for all other parameters. The enzyme ligand-bound site was used as a possible binding site to analyze the potential binding of isolated compounds. Accuracy of results was performed by re-docking the reference ligands. The Discovery Studio 2021 client software allowed us to take the 2D and 3D images.

#### ADME Studies

To analyze the pharmacokinetics of a given molecule that could be used as a drug, ADME studies and licentiousness analyses are needed. The SwissADME web server was used for ADME predictions and drug lawfulness analyses (Daina et al., 2017). Various pharmacokinetic parameters such as physicochemical properties, lipophilicity and water solubility were predicted in this study.

#### *In vitro* intestinal urease activity test

In 96-well plates were incubated at 37°C for 15 min, reaction mixtures consisting of 25 *μ*L of test compounds (each 0.5 mM), 25 *μ*L of enzyme solution (Jack bean urease) and 55 *μ*L of buffers containing 20.79 mM urea. The indophenol method described by Weatherburn (Weatherburn, 1967) was used to determine urease inhibitor activity by measuring ammonia production. Briefly, to each well were added 70 *μ*L of alkaline reagent (0.5% w/v NaOH and 0.1% active chloride, NaOCl) and 45 *μ*L of phenolic reagent (1% w/v phenol and 0.005% w/v sodium nitroprusside). The microplate reader (Molecular Devices, USA) was used to measure the increase in absorbance at 650 nm after 50 min. A final volume of 245 *μ*L was required to perform all reactions in a triplicate. The results (absorbance change per minute) were processed using SoftMax Pro software (Molecular Devices, USA). All assays were performed at pH 6.8. Inhibition percentages were calculated from the formula 100 - (A_test wells_ /A_Control_) × 100. Where “A” is the absorbance of the “test well” as well as the “control”. Thiourea was used as the reference urease inhibitor.

#### *In vitro* intestinal *β*-glucosidase activity test

The method for assessing *β*-glucosidase inhibitory activity described by Ma and collaborator in 2011 with slight modification was used (Ma et al., 2011). A total volume of 180 *μL* was used for inhibition assays in 96-well plates. Distilled water was used to prepare standard solutions of the *β*-glucosidase inhibitors. *β*-glucosidase and *para-nitrophenyl-β-D-* glucuronidase as substrate were prepared in 0.07 M phosphate buffer at pH 7.0. Inhibition assays were performed by adding 20 *μ*L of inhibitor solution to 120 *μ*L of buffer and 20 *μ*L of enzyme solution in 70 mM phosphate buffer (pH 7.0), followed by preincubation at 37°C for 15 minutes. After preincubation, 20 *μ*L of 10 mM *para*-nitrophenyl-*β-D*-glucuronidase as substrate in phosphate buffer was added to each well to start the reaction. The reaction mixture was incubated at 37°C for 15 minutes and then stopped by adding 20 *μ*L of 0.2 M Na_2_CO_3_. 20 *μ*L of DMSO was used as a negative control. Acarbose was used as a positive control. The p-nitrophenol released by *para*-nitrophenyl-*β-D*-glucuronidase at 405 nm was measured to determine *β*-glucosidase activity. The equation below was used to calculate the % inhibition:

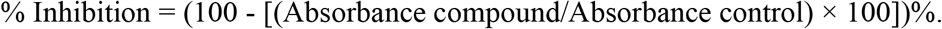

#### Preparation of semi-synthetic derivative (10) from stigmast-4-en-3,6-dione

In a 50 mL flask, 20 mg (4.69 x 10^-5^ mol) of stigmast-4-en-3,6-dione (**10**) were dissolved in 6 mL of methanol and refluxed under stirring until complete dissolution and then allowed to cool (25°C). While under continuous stirring, 2 mg sodium borohydride (NaBH_4_) previously dissolved in 3 mL distilled water was added dropwise and the evolution of the reaction was monitored by TLC until complete disappearance of starting material (20 min). The reaction medium was then transferred to a beaker (250 mL) containing 50 mL of acidified ice-cold distilled water (3 mL of concentrated HCl). After 20 minutes the formation of a precipitate was observed. The medium was then filtered to yield 14.8 mg or 73.3%.

Cell line studies:

## Results and discussion

### Characterization of isolated compounds

Compound **1** obtained as a white powder, exhibited in the HR-ESI-MS (positive-ion mode) a sodium adduct at *m/z* 447.3548 [M+Na]^+^ (calcd. for C_30_H_48_ONa, 447.3597), consistent with the molecular formula of C_30_H_48_O.

All proton and carbon signals of **1** were assigned on the basis of ^1^H NMR, ^1^H-^1^H COSY, HSQC and HMBC experiments. The ^1^H NMR spectrum of compound **1** (Table) exhibited in the upfield region signals of two tertiary methyls resonating at *δ*_H_ 0.53 (Me-18) and 0.81 (Me-19), and four secondary methyls at *δ*_H_ 1.59 (3H; d; *J* = 6.8 Hz; Me-29), 0.98 (3H; d; *J* = 6.9 Hz; Me-27), 0.97 (3H; d; J = 6.5 Hz; Me-26) and at δ_H_ 0.99 (3H; d; *J* = 6.2 Hz; Me-30). Signals of two olefinic protons were also observed at *δ*_H_ 5.17 (1H; dd; *J* = 5.9; 2.3 Hz; H-7) and 5.09 (1H; d; *J* = 6.9 Hz; H-28). In addition, the signals of two methylene protons were observed in this spectrum at *δ*_H_ 4.72 (1H; s; H-21a) and 4.66 (1H; t; *J* = 1.7 Hz; H-21b). The ^13^C NMR spectrum of compound **1** combined to the HMBC spectrum showed a set of 30 carbons. The most interesting resonances were those of olefinic carbons depicted at *δ*_C_ 117.6 (C-6), 139.1 (C-8), 116.4 (C-28), 148.5 (C-24) and those of the methylene carbons at *δ*C 105.8 (C-21), 156.8 (C-20). The backbone of compound **2** was identified as citrostadienol and all the ^1^H and ^13^C NMR spectral data were in good agreement with literature values (Pascal et al., 1993; Schaller, 2010; X. Zhang et al., 2006). Resonances of two methylene protons at *δ*_H_ 4.72 (1H; s; H-21a) and 4.66 (1H; t; *J* = 1.7 Hz; H-21b) giving HSQC correlations with the carbon at δ_C_ 105.8 (C-21) were also observed. The HMBC correlations observed from the protons at *δ*_H_ 0.97 (Me-26) and 1.59 (Me-29) to the carbons at *δ*C 116.4 (C-28), and 148.5 (C-24) as well as the ^1^H-^1^H COSY correlation betweenthe protons at *δ*_H_ 5.17 (H-7) and 2.12 (H-6) allowed us to locate the positions of the double bonds 8 (Figure 2). Thus, the structure of **1** was established as 21-methylene-24-ethylidene lophenol, a previously unreported avenasterol-type phytosterols to which we gave the trivial name Δ^20-21^citrostadienol.

**Figure 2:**
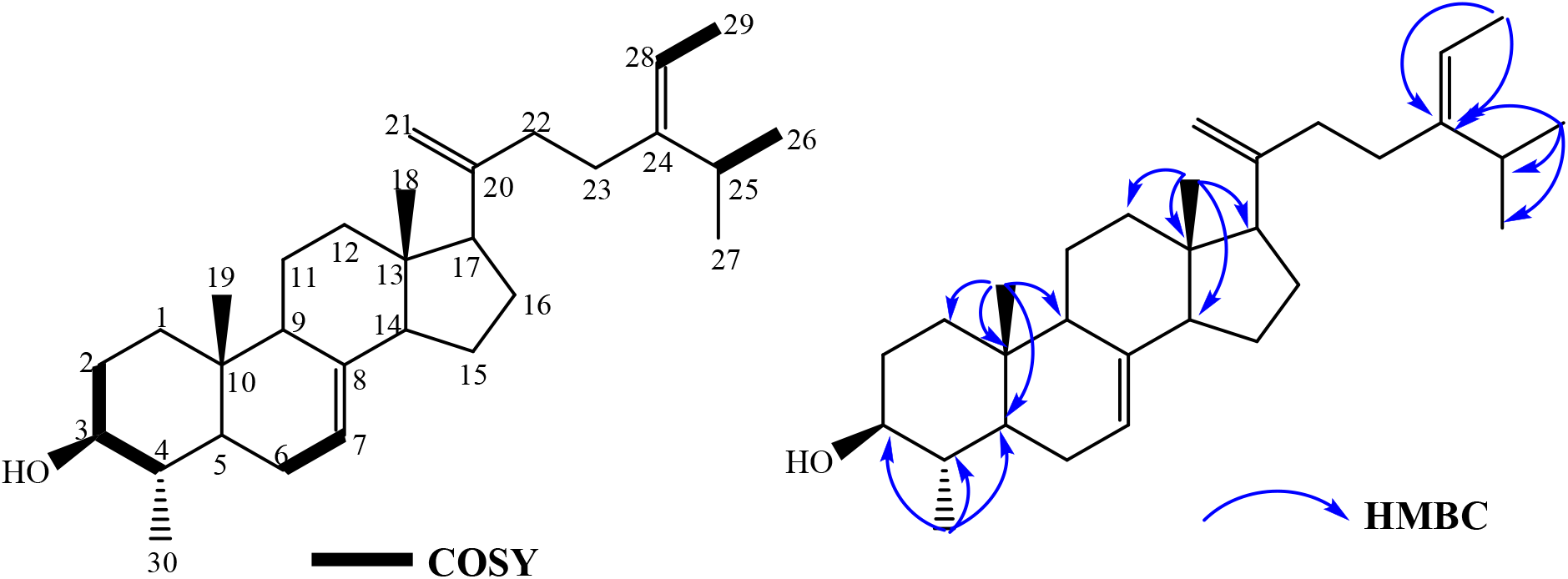
^1^H-^1^H COSY and HMBC correlations of compound **1**

#### **Luteolin (2)** (Cuong et al., 2019)

Yellow powder, ^1^H NMR (600 MHz, Acetone-*d_6_*) *δ* (ppm): 6.59 (1H, s, H-3), 12.99 (1H, s, OH-5), 6.26 (1H, d, *J* = 2.1 Hz, H-6), 6.53 (1H, d, *J* = 2.3 Hz, H-8), 7.47 (1H, dd, *J* = 8.4, 2.2 Hz, H-2’), 7.01 (1H, d, *J* = 8.4 Hz, H-3’), 7.50 (1H, d, *J* = 2.2 Hz, H-6’). ^13^C NMR (150 MHz, Acetone-*d_6_*) *δ* (ppm): 164.3 (C-2), 103.1 (C-3), 182.2 (C-4), 162.3 (C-5), 98.8 (C-6), 164.2 (C-7), 93.8 (C-8), 157.8 (C-9), 104.3 (C-10), 122.6 (C-1’), 119.1 (C-2’), 115.6 (C-3’), 149.4 (C-4’), 145.7 (C-5’), 113.0 (C-6’).

#### **3’,4’,7-trihydroxyflavone (3)** (Yang et al., 2018)

Yellow powder, ^1^H NMR (600 MHz, Acetone-*d_6_*) *δ* (ppm): 6.64 (1H, s, H-3), 7.62 (1H, d, *J* = 8.4 Hz, H-5), 6.79 (1H, dd, *J* = 8.4, 2.0 Hz, H-6), 6.83 (1H, d, *J* = 2.0 Hz, H-8), 7.35 (1H, dd, *J* = 8.3, 2.1 Hz, H-2’), 6.95 (1H, d, *J* = 8.2 Hz, H-3’), 7.58 (1H, d, *J* = 2.1 Hz, H-6’). ^13^C NMR (150 MHz, Acetone-*d_6_*) *δ* (ppm): 146.3 (C-2), 111.3 (C-3), 181.5 (C-4), 125.5 (C-5), 114.0 (C-6), 168.0 (C-7), 98.5 (C-8), 165.7 (C-9), 112.5 (C-10), 117.7 (C-1’), 124.7 (C-2’), 115.6 (C-3’), 147.3 (C-4’), 145.2 (C-5’), 124.5 (C-6’).

#### **Luteolin-5-methyl ether (4)** (Rafieian-Kopaei et al., 2020)

Yellow powder, ^1^H NMR (400 MHz, CD_3_OD) *δ* (ppm): 6.47 (1H, s, H-3), 6.40 (1H, d, *J* = 2.2 Hz, H-6), 6.53 (1H, d, *J* = 2.1 Hz, H-8), 7.33 (1H, dd, *J* = 8.9, 2.3 Hz, H-2’), 6.89 (1H, d, *J* = 8.9 Hz, H-3’), 7.30 (1H, d, *J* = 2.3 Hz, H-6’), 3.88 (3H, s, OCH_3_-5). ^13^C NMR (100 MHz, CD_3_OD) *δ* (ppm): 162.4 (C-2), 105.1 (C-3), 178.8 (C-4), 161.0 (C-5), 96.1 (C-6), 163.6 (C-7), 94.9 (C-8), 159.8 (C-9), 106.8 (C-10), 122.2 (C-1’), 118.5 (C-2’), 115.3 (C-3’), 149.1 (C-4’), 145.6 (C-5’), 112.5 (C-6’), 55.0 (OCH3-5).

#### **Apigenin-8-*C*-*β*-D-glucopyranoside (5)** (Cuong et al., 2015)

Yellow powder, ^1^H NMR (400 MHz, DMSO-*d_6_*) *δ* (ppm): 6.77 (1H, s, H-3), 13.17 (1H, s, OH-5), 6.26 (1H, s, H-6),, 8.02 (2H, d, *J* = 8.02, H-2’/ H-6’), 6.88 (2H, d, *J* = 8.02 Hz, H-3’/ H-5’), 4.67 (1H, dd, *J* = 9.8, 3.4 Hz, H-1’’), 3.82 (1H, dd, *J* = 9.8, 3.4 Hz, H-2’’), 3.25 (1H, m, H-3’’), 3.36 (1H, m, H-4’’), 3.23 (1H, m, H-5’’), 3.76 (1H, d, *J* = 12.1 Hz, H-6a’’), 3.52 (1H, d, *J* = 6.4 Hz, H-6b’’). ^13^C NMR (100 MHz, DMSO-*d_6_*) *δ* (ppm): 164.6 (C-2), 102.9 (C-3), 182.5 (C-4), 161.0 (C-5), 98.6 (C-6), 163.1 (C-7), 105.2 (C-8), 156.7 (C-9), 104.6 (C-10), 122.6 (C-1’), 129.4 (C-2’/C-6’), 116.2 (C-3’/C-5’), 73.9 (C-1’’), 71.2 (C-2’’), 79.1 (C-3’’), 70.9 (C-4’’), 82.3 (C-5’’), 61.7 (C-6’’).

#### **Luteolin-8-*C*-*β*-D-glucopyranoside (Orientin) (6)** (Cuong et al., 2015)

Yellow powder, ^1^H NMR (400 MHz, DMSO-*d_6_*) *δ* (ppm): 6.64 (1H, s, H-3), 13.17 (1H, s, OH-5), 6.25 (1H, s, H-6), 7.52 (1H, dd, *J* = 8.5, 2.2 Hz, H-2’), 6.87 (1H, d, *J* = 8.4 Hz, H-3’), 7.46 (1H, d, *J* = 2.2 Hz, H-6’), 4.67 (1H, dd, *J* = 9.8, 3.4 Hz, H-1’’), 3.83 (1H, m, H-2’’), 3.24 (1H, m, H-3’’), 3.36 (1H, m, H-4’’), 3.28 (1H, m, H-5’’), 3.77 (1H, m, H-6a’’), 3.53 (1H, m, H-6b’’). ^13^C NMR (100 MHz, DMSO-*d_6_*) *δ* (ppm): 164.6 (C-2), 102.8 (C-3), 182.4 (C-4), 160.7 (C-5), 98.6 (C-6), 162.9 (C-7), 104.9 (C-8), 156.4 (C-9), 104.4 (C-10), 122.3 (C-1’), 119.8 (C-2’), 116.0 (C-3’), 150.1 (C-4’), 146.2 (C-5’), 114.4 (C-6’), 73.8 (C-1’’), 71.2 (C-2’’), 79.1 (C-3’’), 71.0 (C-4’’), 82.3 (C-5’’), 61.7 (C-6’’).

#### **Luteolin-6-*C*-*β*-D-glucopyranoside (Homoorientin) (7)** (Cuong et al., 2015)

Yellow powder, ^1^H NMR (400 MHz, DMSO-*d_6_*) *δ* (ppm): 6.70 (1H, s, H-3), 13.58 (1H, s, OH-5), 6.50 (1H, s, H-8), 7.44 (1H, dd, *J* = 8.2, 2.3 Hz, H-2’), 6.87 (1H, d, *J* = 8.3 Hz, H-3’), 7.42 (1H, d, *J* = 2.3 Hz, H-6’), 4.60 (1H, d, *J* = 9.8 Hz, H-1’’), 4.07 (1H, d, *J* = 9.2 Hz, H-1’’), 3.21 (1H, m, H-3’’), 3.14 (1H, m, H-4’’), 3.18 (1H, m, H-5’’), 3.70 (1H, m, H-6a’’), 3.42 (1H, m, H-6b’’). ^13^C NMR (100 MHz, DMSO-*d_6_*) δ (ppm): 164.0 (C-2), 103.7 (C-3), 182.3 (C-4), 161.2 (C-5), 109.3 (C-6), 163.7 (C-7), 93.9 (C-8), 156.7 (C-9), 103.9 (C-10), 121.9 (C-1’), 116.4 (C-2’), 116.0 (C-3’), 150.1 (C-4’), 146.2 (C-5’), 114.4 (C-6’), 73.4 (C-1’’), 70.6 (C-2’’), 79.4 (C-3’’), 71.0 (C-4’’), 82.0 (C-5’’), 61.7 (C-6’’).

#### **Butin (8)** (Tian et al., 2004)

A orange powder, ^1^H NMR (600 MHz, Acetone-*d_6_*) *δ* (ppm): 5.40 (1H, dd, *J* = 12.8, 2.9 Hz, H-2), 3.02 (1H, dd, *J* = 16.7, 12.8 Hz, H-3a), 2.68 (1H, dd, *J* = 16.7, 3.0 Hz, H-3b), 7.73 (1H, d, *J* = 8.6 Hz, H-5), 6.58 (1H, dd, *J* = 8.6, 2.3 Hz, H-6), 6.43 (1H, d, *J* = 2.3 Hz, H-8), 7.05 (1H, d, *J* = 1.9 Hz, H-2’), 6.87 (1H, d, *J* = 8.1 Hz, H-5’), 6.89 (1H, dd, *J* = 8.2, 1.9 Hz, H-6’). ^13^C NMR (150 MHz, Acetone-*d_6_*) δ (ppm): 79.6 (C-2), 43.8 (C-3), 189.6 (C-4), 128.5 (C-5), 110.2 (C-6), 163.5 (C-7), 102.7 (C-8), 163.2 (C-9), 114.3 (C-10), 131.2 (C-1’), 113.7 (C-2’), 145.6 (C-3’), 145.6 (C-4’), 114.7 (C-5’), 118.2 (C-6’).

#### **Stigmast-4-en-3,6-dione (9)** (Greca et al., 1990)

White powder, ^1^H NMR (600 MHz, C_5_D_5_N) *δ* (ppm): 1.69 (1H, m, H-1a), 1.52 (1H, m, H-1b), 1.78 (1H, m, H-2a), 1.39 (1H, m, H-2b), 6.41 (1H, s, H-4), 2.67 (1H, dd, *J* = 16.1, 4.4 Hz, H-7a), 2.05 (1H, dd, *J* = 16.1, 12.3 Hz, H-7b), 1,78 (1H, m, H-8), 1.24 (1H, m, H-9), 1.48 (2H, m, H-11), 2.00 (1H, m, H-12a), 1,15 (1H, m, H-12b), 1.02 (1H, m, H-14), 1.51 (1H, m, H-15a), 1.31 (1H, m, H-15b), 1.83 (2H, m, H-16), 1.03 (1H, m, H-17), 0.68 (3H, s, H-18), 1.00 (3H, s, H-19), 1.38 (1H, m, H-20), 0.98 (3H, d, *J* = 6.3 Hz, H-21), 2.39 (1H, m, H-22a), 1,05 (1H, m, H-22b), 1.24 (2H, m, H-23), 1.49 (1H, m, H-24), 1.68 (1H, m, H-25), 0.88 (3H, d, *J* = 6.1 Hz, H-26), 0.85 (3H, d, *J* = 6.8 Hz, H-27), 1.31 (2H, m, H-28), 0.90 (3H, t, H-29). ^13^C NMR (150 MHz, C_5_D_5_N) *δ* (ppm): 35.1 (C-1), 33.9 (C-2), 198.9 (C-3), 125.3 (C-4), 160.8 (C-5), 201.6 (C-6), 46.5 (C-7), 39.1 (C-8), 50.4 (C-9), 34.0 (C-10), 20.7 (C-11), 39.5 (C-12), 42.4 (C-13), 55.8 (C-14), 23.1 (C-15), 28.1 (C-16), 56.2 (C-17), 11.7 (C-18), 16.9 (C-19), 36.0 (C-20), 18.6 (C-21), 33.8 (C-22), 26.1 (C-23), 45.8 (C-24), 29.2 (C-25), 19.3 (C-26), 18.9 (C-27), 23.1 (C-28), 11.9 (C-29).

#### **Stigmast-4-en-3*β*,6α-diol (10)** (Zhao et al., 2005)

White powder, ^1^H NMR (400 MHz, Acetone-*d_6_*) *δ* (ppm): 1.73 (1H, m, H-1a), 1.69 (1H, m, H-1b), 1.40 (1H, m, H-2a), 1.27 (1H, m, H-2b), 4.08 (1H, m, H-3), 5.74 (1H, d, *J* = 1.7 Hz, H-4), 4.11 (1H, m, H-6), 2.02 (2H, m, H-7), 1,30 (1H, m, H-8), 0.72 (1H, m, H-9), 1.50 (1H, m, H-11a), 1.38 (1H, m, H-11b), 2.05 (1H, m, H-12a), 1,16 (1H, m, H-12b), 1.17 (1H, m, H-14), 1.23 (2H, m, H-15), 1.69 (2H, m, H-16), 1.05 (1H, m, H-17), 1.05 (3H, s, H-18), 0.74 (3H, s, H-19), 1.42 (1H, m, H-20), 0.96 (3H, d, *J* = 6.4 Hz, H-21), 1.67 (1H, m, H-22a), 1,40 (1H, m, H-22b), 1.92 (2H, m, H-23), 0.98 (1H, m, H-24), 1.70 (1H, m, H-25), 0.88 (3H, m, H-26), 0.84 (3H, m, H-27), 1.30 (2H, m, H-28), 0.88 (3H, m, H-29). ^13^C NMR (100 MHz, Acetone*d_6_*) *δ* (ppm): 36.4 (C-1), 34.3 (C-2), 67.4 (C-3), 121.5 (C-4), 147.5 (C-5), 67.0 (C-6), 42.5 (C-7), 29.5 (C-8), 54.5 (C-9), 37.4 (C-10), 20.8 (C-11), 39.7 (C-12), 42.4 (C-13), 56.1 (C-14), 25.8 (C-15), 24.0 (C-16), 56.0 (C-17), 19.2 (C-18), 11.4 (C-19), 36.0 (C-20), 18.2 (C-21), 33.7 (C-22), 28.0 (C-23), 45.8 (C-24), 29.1 (C-25), 19.2 (C-26), 18.4 (C-27), 22.8 (C-28), 11.3 (C-29).

#### **Emodin (11)** (Ko et al., 1995)

Orange powder, ^1^H NMR (600 MHz, Acetone-*d_6_*) δ (ppm): 12.10 (1H, s, OH-1), 7.15 (1H, s, H-2), 7.58 (1H, d, *J* = 1.6 Hz, H-4), 7.26 (1H, d, *J* = 2.4 Hz, H-5), 6.67 (1H, d, *J* = 2.4 Hz, H-7), 12.20 (1H, s, OH-8), 2.48 (3H, s, H-1’). ^13^C NMR (150 MHz, Acetone-*d_6_*) δ (ppm): 165.4 (C-1), 124.0 (C-2), 148.6 (C-3), 120.5 (C-4), 133.3 (C-4a), 108.9 (C-5), 162.3 (C-6), 107.9 (C-7), 165.9 (C-8), 109.1 (C-8a), 190.7 (C-9), 113.6 (C-9a), 181.4 (C-10), 135.6 (C-10a), 21.0 (C-1’).

#### **Chrysophanol (12)** (Prateeksha et al., 2019)

Orange powder, ^1^H NMR (600 MHz, CDCl3) δ (ppm): 12.10 (1H, s, OH-1), 7.12 (1H, m, H-2), 7.67 (1H, d, *J* = 1.8 Hz, H-4), 7.84 (1H, dd, *J* = 7.5, 1.2 Hz, H-5), 7.69 (1H, d, *J* = 8.0 Hz, H-6), 7.31 (1H, dd, *J* = 8.5, 1.2 Hz, H-7), 12.03 (1H, s, OH-8), 2.49 (3H, s, H-1’). ^13^C NMR (150 MHz, CDCl_3_) δ (ppm): 162.4 (C-1), 124.5 (C-2), 149.3 (C-3), 121.3 (C-4), 133.2 (C-4a), 119.9 (C-5), 136.9 (C-6), 124.3 (C-7), 162.7 (C-8), 115.8 (C-8a), 192.5 (C-9), 113.7 (C-9a), 182.0 (C-10), 133.6 (C-10a), 22.2 (C-1’).

#### **Physcion (**13) (Ko et al., 1995)

Orange powder, ^1^H NMR (400 MHz, CDCl_3_) δ (ppm): 12.13 (1H, s, OH-1), 7.09 (1H, d, *J* = 1.5 Hz, H-2), 7.63 (1H, d, *J* = 1.7 Hz, H-4), 7.37 (1H, d, *J* = 2.6 Hz, H-5), 6.69 (1H, d, *J* = 2.5 Hz, H-7), 12.33 (1H, s, OH-8), 2.46 (3H, s, H-1’), 3.94 (3H, s, H-1’’). ^13^C NMR (100 MHz, CDCl_3_) *δ* (ppm): 162.5 (C-1), 124.5 (C-2), 148.4 (C-3), 121.3 (C-4), 133.2 (C-4a), 108.2 (C-5), 166.5 (C-6), 106.7 (C-7), 165.2 (C-8), 110.2 (C-8a), 190.8 (C-9), 113.7 (C-9a), 181.8 (C-10), 135.2 (C-10a), 22.1 (C-1’), 56.1 (C-1’’).

### Molecular Docking Studies

To assess the tendency of compounds isolated from *C. mimosides*, shown in Figure 1, to bind to the urease and *β*-glucosidase enzyme, we conducted a series of in silico experiments based primarily on molecular docking and molecular dynamics (Table 2 and 3). The protein (PDB ID: 3LA4 and 2XHY) and ligands prepared as described in Materials and Methods were docked into the active site of the *β*-glucosidase and urease enzyme. The result of this calculation is shown in Table 2 and 3, while the docking results are shown in Figure 3, 4, 5 and 6.

**Table 1:** ^13^C NMR (100 MHz) and ^1^H NMR (400 MHz) data of compound **1** (CDCl_3_): *δ* in ppm, J in Hz

| Positions | $\delta^{13}\text{C}$ | $\delta^1\text{H}$ (mult, J) |
| --- | --- | --- |
| **1** | 36.9 | 1.84 (m, 1H)<br>1.81 (m, 1H) |
| **2** | 30.9 | 1.80 (m, 1H)<br>1.46 (m, 1H) |
| **3** | 76.2 | 3.12 (m, 1H) |
| **4** | 40.2 | 1.30 (m, 1H) |
| **5** | 46.6 | 1.01 (m, 1H) |
| **6** | 26.6 | 2.13 (m, 1H)<br>2.09 (m, 1H) |
| **7** | 117.6 | 5.17(dt, $J = 5.9, 2.3$ Hz, 1H) |
| **8** | 139.1 | - |
| **9** | 49.6 | 1.64 (m, 1H) |
| **10** | 34.8 | - |
| **11** | 21.3 | 1.55 (m, 1H)<br>1.45 (m, 1H) |
| **12** | 39.4 | 2.02 (m, 1H)<br>1.22 (m, 1H) |
| **13** | 43.3 | - |
| **14** | 54.9 | 1.80 (m, 1H) |
| **15** | 22.8 | 1.51 (m, 1H)<br>1.25 (m, 1H) |
| **16** | 29.6 | 1.25 (m, 1H) |
| **17** | 55.9 | 1.22 (m, 1H) |
| **18** | 11.8 | 0.53 (s, 3H) |
| **19** | 14.1 | 0.83 (s, 3H) |
| **20** | 156.8 | - |
| **21** | 105.8 | 4.72 (s, 1H)<br>4.66 (t, $J = 1.7$ Hz) |
| **22** | 29.6 | 1.25 (m, 1H) |
| **23** | 27.9 | 1.91 (m, 1H)<br>1.27 (m, 1H) |
| **24** | 145.8 | - |
| **25** | 28.5 | 2.85 (m, 1H) |
| **26** | 21.0 | 0.97 (d, $J = 6.5$ Hz, 3H) |
| **27** | 20.9 | 0.98 (d, $J = 6.9$ Hz, 3H) |
| **28** | 116.4 | 5.09 (q, $J = 6.9$ Hz, 1H) |
| **29** | 12.7 | 1.59 (d, $J = 6.8$ Hz, 3H) |
| **30** | 15.1 | 0.99 (d, $J = 6.2$ Hz, 3H) |

**Table 2:** Binding score and RMSD from *β*-glucosidase

| Compound Name | Binding score (kcal/mol) | RMSD (Å) |
| --- | --- | --- |
| Luteolin-8-*C*-β-D-glucopyranoside (6) | -7.33 | 1.87 |
| Nonadecanoic acid (14) | -7.23 | 1.33 |
| β-sitosterol 3-*O*-β-D-glucopyranoside (16) | -7.21 | 2.91 |
| Luteolin-6-*C*-β-D-glucopyranoside (7) | -7.19 | 2.06 |
| β-sitosterol (15) | -6.98 | 1.28 |
| Apigenin-8-*C*-β-D-glucopyranoside (5) | -6.82 | 1.08 |
| 21-methylene-24-ethylidene lophenol (1) | -6.70 | 1.60 |
| Stigmast-4-en-3β,6α-diol (10) | -6.57 | 2.48 |
| Stigmat-4-en-3,6-dione(9) | -6.56 | 1.73 |
| Butin (8) | -6.03 | 1.33 |
| Luteolin (2) | -5.97 | 4.31 |
| Emodin (11) | -5.71 | 2.11 |
| Luteolin-5-methyl ether (4) | -5.45 | 3.55 |
| Physcion (13) | -5.25 | 1.95 |
| 3',4',7-trihydroxyflavone (3) | -5.01 | 2.20 |
| Chrysophanol (12) | -4.02 | 2.95 |
| Reference (Acarbose) | -7.51 | 2.03 |

**Table 3:** Binding score and RMSD from urease

| Compound Name | Binding score (kcal/mol) | RMSD (Å) |
| --- | --- | --- |
| ***β-sitosterol 3-O-β-D-glucopyranoside (16)*** | -7.42 | 1.74 |
| ***Lutéolin-8-C-β-D-glucopyranoside (6)*** | -7.27 | 1.08 |
| ***21-methylene-24-ethylidene lophenol (1)*** | -7.23 | 1.46 |
| ***Apigenin-8-C-β-D-glucopyranoside (5)*** | -7.20 | 0.63 |
| ***Stigmast-4-en-3β,6α-diol (10)*** | -7.05 | 1.72 |
| ***Luteolin-6-C-β-D-glucopyranoside(7)*** | -6.95 | 1.60 |
| ***Stigmat-4-en-3,6-dione (9)*** | -6.80 | 1.36 |
| ***β-sitosterol (15)*** | -6.57 | 1.76 |
| ***Nonadecanoic acid (14)*** | -6.41 | 1.21 |
| ***Physine (13)*** | -6.18 | 0.79 |
| ***Luteolin-5-methyl ether (4)*** | -5.82 | 2.56 |
| ***Emodin (11)*** | -5.36 | 3.27 |
| ***Luteolin (2)*** | -5.23 | 3.05 |
| ***Chrysophanol (12)*** | -5.21 | 2.41 |
| ***3',4',7-trihydroxyflavone (3)*** | -5.18 | 1.69 |
| ***Butin (8)*** | -4.90 | 2.04 |
| ***Reference (thiourea)*** | -3.13 | 2.15 |

**Figure 3:**
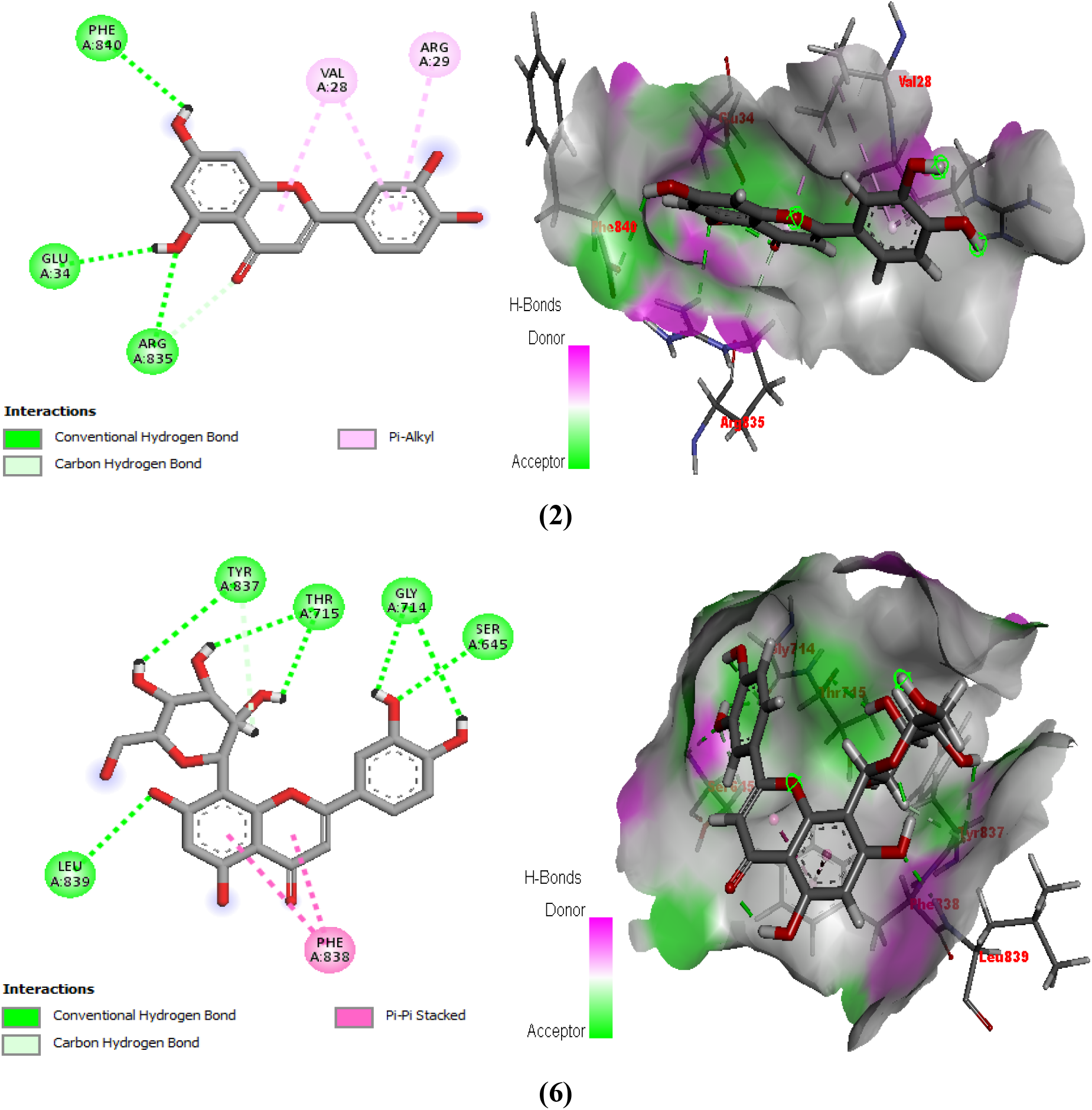
2D and 3D view of binding interaction of urease

**Figure 4:**
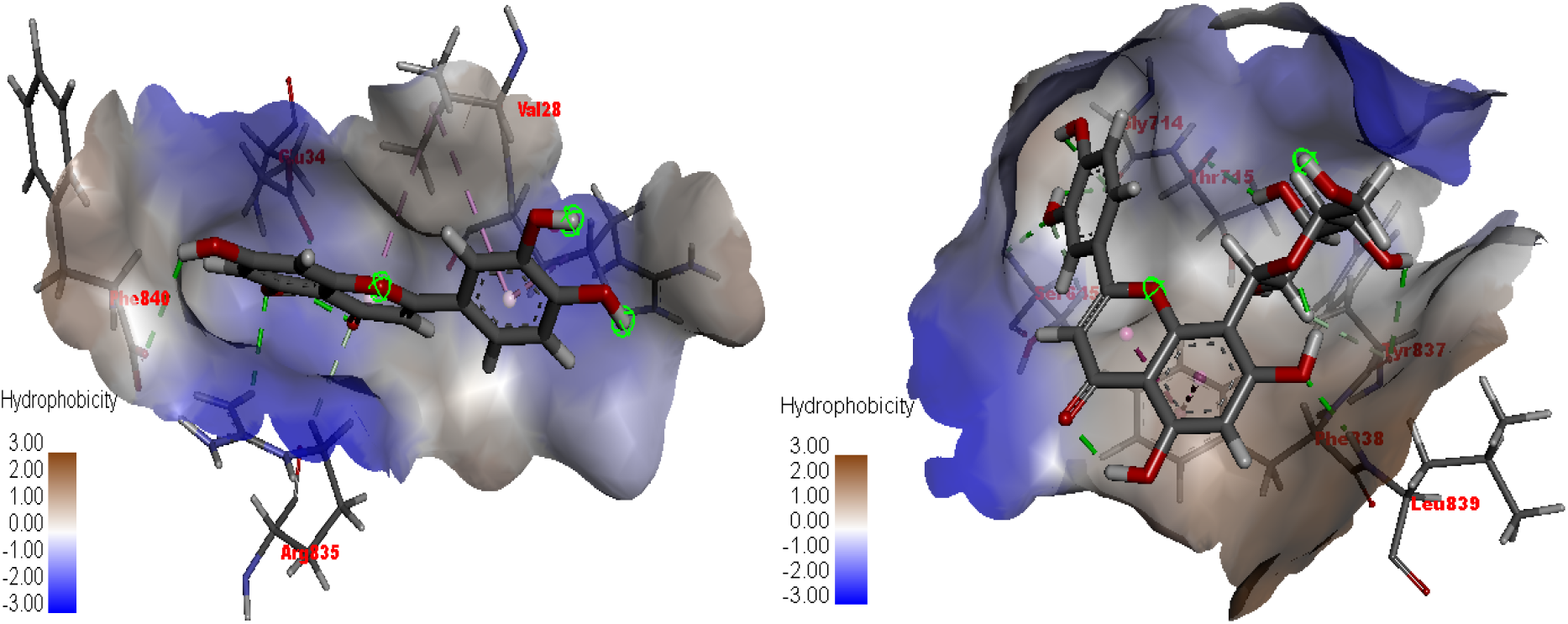
3D view interaction hydrophobicity of compounds **2** and **6**

**Figure 5:**
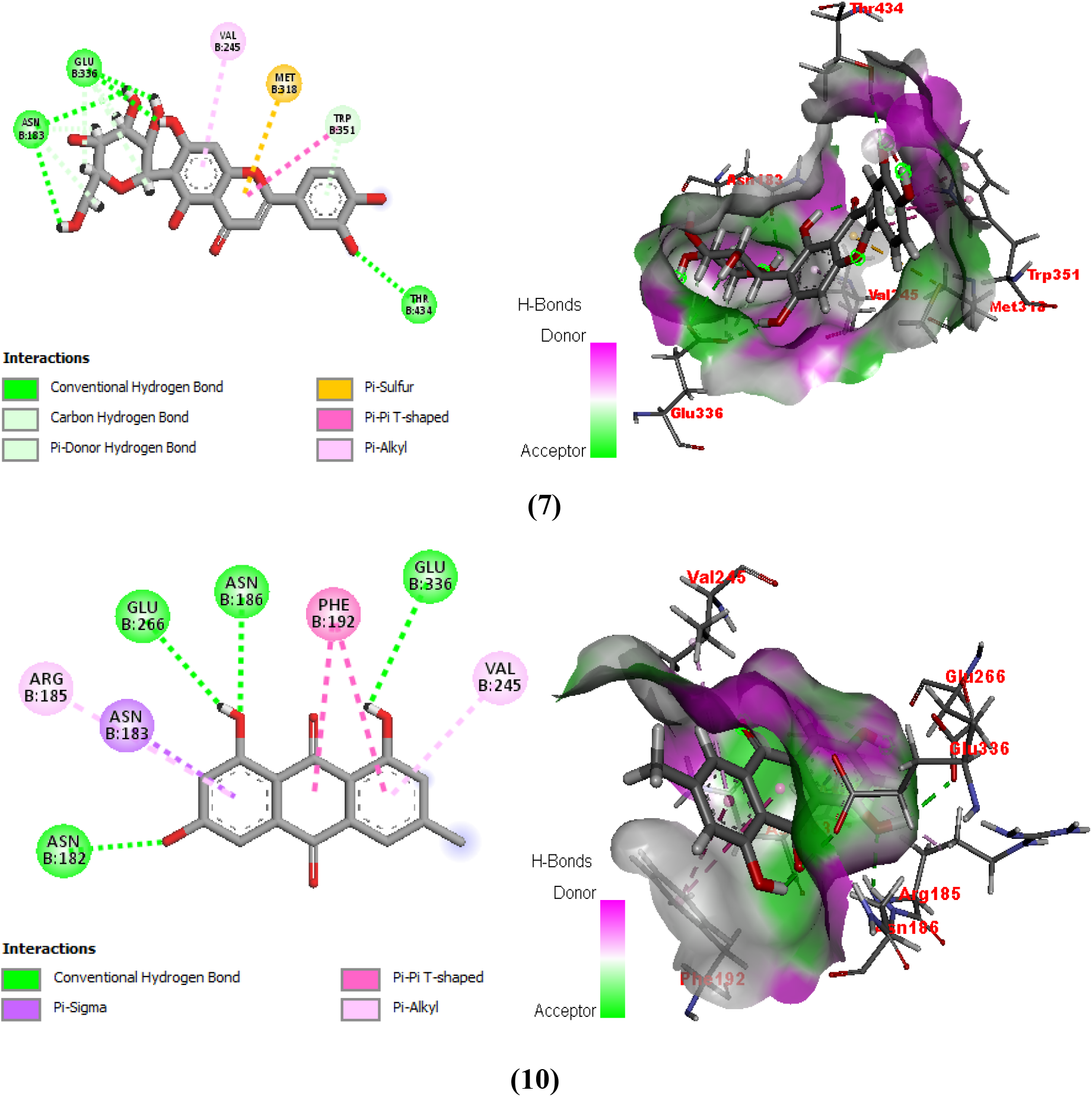
2D and 3D view of binding interaction of *β*-glucosidase

**Figure 6:**
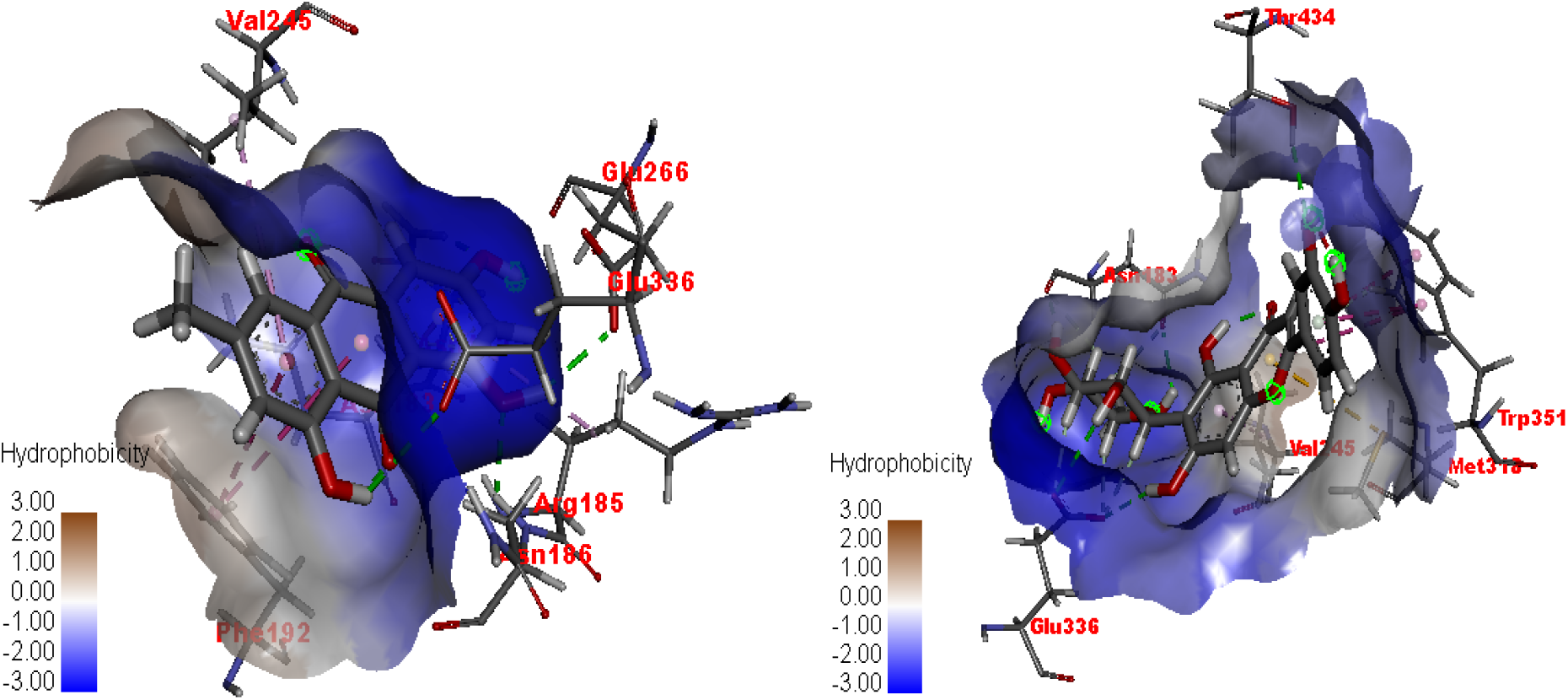
3D view interaction hydrophobicity of compounds **10** and **7**

### In silico analysis of urease

The docking results showed that compounds **16**, **6**, **5**, **1**, and **7** had the highest binding affinities of −7.42, 7.27, 7.20, 7, 11, and 6.95 kcal/mol, respectively. On the other hand, compound 4 showed the lowest binding affinity of −4.90 kcal/mol compared to the other compounds. Hydrogen bonds were observed in the majority of protein-ligand complexes but not in those involving compounds **1**, **8**, **9**, and **13** on some residues, namely Ala80, Asp295, Asp730, Glu34, Glu742, Gly714, Gly641, Ile148, Phe838, Ser421, and Tyr309. Based on the binding interactions, one critical residue (Glu742) of the 3LA4 protein interacted only with compounds **4**, **7**, **12**, and **16**, through hydrogen or hydrophobic interactions.

However 66.66% of the compounds studied showed better docking protocol accuracy than the reference compound with RSMDs between 0.63 and 1.79 Å.

### In silico analysis of *β*-glucosidase

The molecular docking studies showed the best binding interaction pattern of the most important compounds, **6, 14, 16, 7, 15, 5, 1** and **10**, inside the active pocket of the protein, with a binding score of −7.3, −7.23, 7.21, 7.1 −6.9, −6.8, −6.7, and −6.5 kcal/mol, respectively. In addition the reference compound exhibited the best binding interaction pattern within the active pocket of the protein with a binding score of 7.5 kcal/mol. However 50 % of the compounds (**6, 14, 15, 5, 1, 9, 8** and **13**) studied showed better docking protocol accuracy than the reference compound with RSMDs between 1.08 and 1.95.

Considering the recovered binding mode, we observed that the compound **7** (Table 2 and Figure 3A) covered and interacted with almost all regions of the *β*-glucosidase binding site. In fact, the cathecol function of ring B made polar contact with the Glu336 backbone while the glucopyraonside unit bound to ring A made polar contact with the Glu180 backbone, thus establishing a series of H-bonds with Glu180 (2.97 Å), Glu180 (2.89 Å), Glu336 (2.66 Å) and Glu336 (2.68 Å). Moreover this compound also established a series of van der Waal interactions with almost all the amino acid residues and the most important of these amino acids were Trp351, Phe192, Tyr317, Phe433, Val245, Asn186 and Glu336. The compound **11**, although having a low score with the protein, showed a better interaction pattern with amino acids such as Asn183 (4.41 Å) Asn183 (3.94 Å) and Glu266 (3.16 Å). This could be explained on the one hand by the presence of the quinone group and on the other hand by the presence of an intramolecular hydrogen bond. Emodin also established van der Waal interactions with Asn183, Phe192, Val245 and Glu336.

On the other hand the result of this calculation confirms that amino acids such as phenylamine, tryptophan and methionine were the active participants in the formation of van der Waal interactions for all the compounds studied. We could also observe with the program discovery studio visualizer, that all the studied compounds presented bonds of hydrophobicity going from −3 to 3 (figure 4 and 6).

### *In Vitro* Analysis: urease Inhibition Potential

The isolated compounds were studied for their inhibition potential against urease using a SpectraMax M2 reader for 96-well microplates. All isolated compounds inhibited the urease enzyme with inhibition percentages ranging from 53.42 to 83.05 % (Figure 7), except for compounds **1** and **16**.

**Figure 7:**
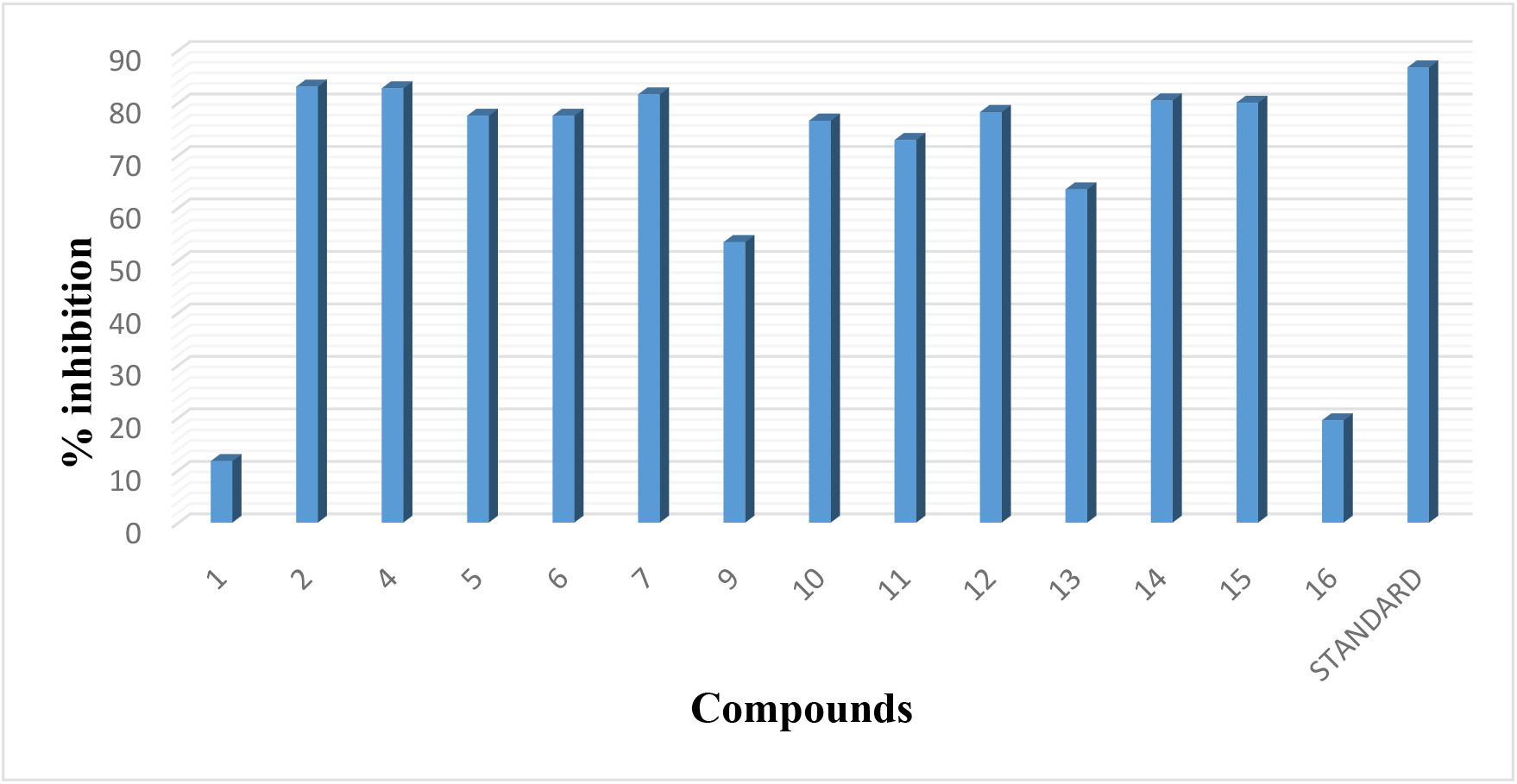
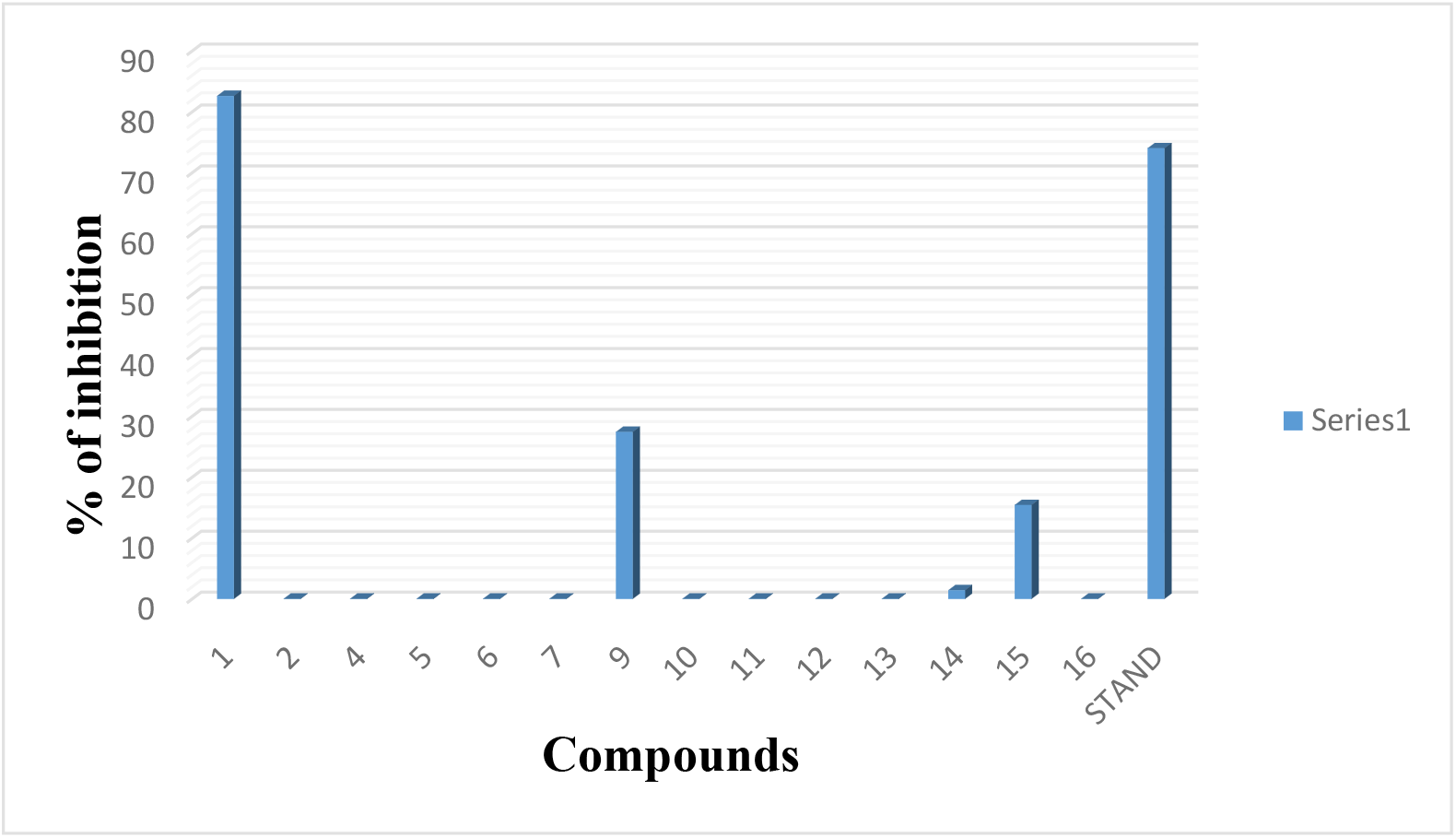
Graphical representation of % inhibition of all compounds (**1**, **2**, **5**-**7**, **9**-**16**) compared against the standard.

Data obtained from the *in vitro* urease inhibition assay demonstrated that the type of secondary metabolite and its substitution pattern are important in determining the structure-activity relationship against urease. The most active compound is compound **2**, whose activity can be attributed to the cathecol function of the flavonoid B-ring and the presence of the hydroxyl groups at positions 5 and 7 of the A-ring. Moreover, the presence of a glucose unit in position C-8 of the A ring decreases the inhibitory activity of the latter compared to that fixed in position C-6 of the same ring. In the case of anthraquinones, the presence of a hydroxyl or a methoxyl in position 6 of the A ring decreases the inhibitory power of the latter. In the case of steroids, the inhibitory power depends on the nature of the steroid skeleton and the different substitutions found within it. The most obvious case is that of sterol (compound **15** and **16**) where the substitution of hydrogen by glucose unit considerably reduces its inhibitory power towards the urease enzyme. In addition to these observations, the reduction of the ketone function of compound **9** to alcohol (Compound **10**) considerably increased its inhibitory power.

In view of their inhibition percentages, we then determined their inhibitory concentrations. It appears that the evaluated compounds presented inhibition concentrations ranging from 1.224±0.43 to 6.678±0.11 *μ*M (Figure 8), whereby compounds **7**, **2** and **14** presented the best inhibitory concentrations. In addition, all the evaluated compounds exhibited higher inhibitory concentration than the reference compound.

**Table 4:** In vitro study – % enzyme inhibition and IC_50_ of isolated compounds

| Compounds | % inhibition | $\text{IC}_{50} \pm \text{SEM}$<br>($\mu\text{M}$) |
| --- | --- | --- |
| **7** | 81.57 | $1.224 \pm 0.43$ |
| **2** | 83.05 | $1.303 \pm 0.28$ |
| **14** | 80.42 | $1.946 \pm 0.31$ |
| **15** | 80.00 | $2.783 \pm 0.24$ |
| **4** | 82.73 | $3.869 \pm 0.24$ |
| **5** | 77.52 | $4.730 \pm 0.22$ |
| **11** | 72.89 | $5.196 \pm 0.14$ |
| **12** | 78.21 | $5.915 \pm 0.25$ |
| **10** | 76.57 | $6.678 \pm 0.11$ |
| **1** | 11.70 | - |
| **6** | 77.52 | - |
| **9** | 53.42 | - |
| **13** | 63.47 | - |
| **16** | 19.47 | - |
| **Standard (Thiourea)** | 86.73 | 18.61±0.11 |

### *In vitro* analysis: *β*-glucosidase inhibition potential

In the present study, the inhibitory activity of *β*-glucosidase was determined against the enzyme obtained from *Escherichia coli*. Acarbose was used as the standard. The results clearly suggest that flavonoids and anthraquinones obtained from the aerial part of *Cassia mimosoides*, did not inhibit *β*-glucosidase (Figure 1).

In the *β*-glucosidase inhibition assay, only an isolated new steroid, namely compound **1** (21-methylene-24-ethylidene lophenol) strongly inhibited *β*-glucosidase compared with the standard acarbose (Figure 9). Furthermore, compound **1** showed the strongest inhibitory activity when compared against all other isolated compounds and acarbose. Our results obviously demonstrate that the isolated flavonoids did not inhibit *β*-glucosidase, this could be due to the orientation of the beta bond of the glucosidase enzyme as the work of Ahmed and collaborator in 2014 highlighted the inhibitory potency of flavonoids isolated from *Albizzia Lebbeck* Benth towards α-glucosidase (Ahmed et al., 2014).

### Drug similarity analyses and ADME studies

Similarity analyses with reference compounds and ADME predictions for the natural compounds studied were performed via the SwissADME web server (Daina et al., 2017). The results are presented in Tables 1S and 2S. Compliance analyses to reference compounds and ADME predictions were also performed for comparison. In these analyses, no secondary metabolites studied had more than twelve hydrogen bond acceptors. However, compounds 5, 6 and 7 have the highest number of hydrogen bond acceptors with values between ten and eleven. On the other hand, although compounds **5**, **6** and **7** obey the Lipinski rule while possessing more than five hydrogen bond donors. The molecular masses of the natural compounds studied are between 254.24 and 576.85 g/mol.

According to Veber’s rule and Muegge’s rule, the rotational bonds of a molecule should not be greater than **10** and **15**, respectively. The results show that, the studied molecules have a maximum of nine rotational bonds and obey the different rules, except for compound 14 which had seventeen rotational bonds. This can be explained by the absence of double bonds and ring in its structure.

The aliphatic degree and solubility of a molecule are predicted by the fraction of sp3 carbon atoms. Furthermore, the clinical success rate of a given molecule is characterized by the increase in saturation rate (Lovering et al., 2009; Wei et al., 2020). In this study, the fraction of sp3 carbon atoms was found to be greater than 0.25 in most par of the compounds and five of the compounds (compounds **1, 14, 15, 9**, and **16**) exhibited much higher degrees of saturation. The molar refractivity of the studied compounds was also predicted in the range of 68.76 −137.56, and it was observed that all the molecules obeyed Ghose’s rule.

The drug character of a given molecule was predicted using topological polar surface area (TPSA) (Wei et al., 2020). According to Veber’s rule, the polar surface area should not be greater than 140 Å2. From these results, it can be seen that, the polar surface area of the majority of the studied compounds are less than 111.13 Å2. However, compounds 5, 6 and 7 showed a polar surface area greater than the reference drug (Acarbose). According to Ghose’s rule, the average partition coefficient of a given molecule, which is the average of iLOGP (Daina et al., 2014), XLOGP3 (Chinese Academy of Sciences, 2007), WLOGP (Wildman & Crippen, 1999), MLOGP (Lipinski et al., 2001) and SILICOS-IT (Ali et al., 2012), should be between −0.4 and 5.6. In this study, this coefficient is between −0.41 and 2.38. However, compounds 1, 14, 15, and 9 have coefficients higher than the standard (6.28-7.25); this could be due to their low solubility.

The water solubility of the studied secondary metabolites (ESOL, ALI and SILICOS-IT (Ali et al., 2012)) varies from poorly soluble to soluble. The high value of gastrointestinal absorption of compound 12 may therefore imply that it can cross the blood-brain barrier. We were also able to observe a variation in skin permeability −1.91 to −9.14 cm/s (Potts & Guy, 1992). These results also show that all the molecules obeyed the rules of Lipinski (86.66%), Ghose (53.33%), Veber (73.33%), Egan (53.33%) and Muegge (46.66%), and Abbott’s bioavailability scores (Martin, 2005)were predicted at 0.55 for 80% of the molecules studied.

In the PAINS analyses, 40% of the studied compounds did not show an alert, while 60% showed an alert due to the presence of catechol A and quinone A groups (Baell & Holloway, 2010). In Brenk’s analyses (Brenk et al., 2008), no alert was observed for 40% of the compounds studied. However, the presence of an isolated alkene function and a catechol function showed a 60% alert. According to the likelihood analyses described by Teague and collaborator (Teague et al., 1999), 60% of the studied compounds presented violations on several parameters (molecular weight, XLOGP3). Finally, the synthetic accessibility scores were obtained within a range of 2.47 to 8.02.

## Conclusion

The genus *Cassia* is a significant source of secondary metabolites that are physiologically active and come from several chemical classes. The current research discusses the spectroscopic elucidation of the structure and enzymatic activity in-silico and in-vitro of fifteen known compounds as well as a new unidentified avenasterol derivative called 21-methylene-24-ethylidene lophenol. For the first time, the inhibitory effects of these compounds on urease and *β*-glucosidase were investigated, and molecular docking studies were also carried out to confirm the structure-activity relationship. The inhibitory activity of all the substances tested against urease was higher (1.224±0.43 IC_50_ > 6.678±0.11 M) than that of thiourea (IC_50_ = 18.61±0.11 M). According to the results of the molecular docking, compound **7** significantly inhibits urease by forming a stable ligand-urease complex through hydrogen bonding, van der Waal, and hydrophobic interactions. Our results from in-vitro experimentation and molecular docking studies demonstrated that flavonoids, anthraquinones, fatty acid and steroid isolated from the aerial part of *Cassia mimosoïdes* significantly inhibit urease and *β*-glucosidase enzymes. These results suggest that the activity of this plant may be due to the synergistic effect of active compounds, including those investigated in the present studies; therefore, this plant is a potential candidate for obtaining new remedies against infectious diseases caused by urease-producing bacteria

## Acknowledgements

The authors would like to thank CUI-TWAS-UNESCO and the COMSATS University Islamabad, Abbottabad campus for providing laboratory facilities.

## Notes

### Competing Interest Statement

The authors have declared no competing interest.

